# Effects of IgG and IgM autoantibodies on non-infected erythrocytes is related to ABO blood group in *Plasmodium vivax* malaria and is associated with anemia

**DOI:** 10.1101/715110

**Authors:** Luiza Carvalho Mourão, Camila Maia Pantuzzo Medeiros, Gustavo Pereira Cardoso-Oliveira, Paula Magda da Silva Roma, Jamila da Silva Sultane Aboobacar, Beatriz Carolina Medeiros Rodrigues, Ubirajara Agero, Cor Jesus Fernandes Fontes, Érika Martins Braga

## Abstract

Autoantibodies play an important role in the destruction of non-infected red blood cells (nRBCs) during malaria. However, the relationship between this clearance and ABO blood groups is yet to be fully enlightened, especially for *Plasmodium vivax* infections. Here we show that anti-RBC IgG and IgM are increased in anemic patients with acute vivax malaria. Furthermore, both antibodies are able to decrease the deformability of nRBCs, but only IgG can induce *in vitro* erythrophagocytosis. Such effects are enhanced in type O erythrocytes, suggesting that individuals from this blood group infected with *P. vivax* malaria may be more susceptible to develop anemia.

## 1. Introduction

*Plasmodium vivax* is the most widely distributed species that cause malaria in humans, accounting for approximately 74.1% of the cases registered in the Americas in 2017 [1]. One of the major complications of *P. vivax* infection is anemia [2, 3], which is thought to arise from the destruction of both infected and non-infected RBCs [4]. Recently we have demonstrated that autoantibodies from *P. vivax*-infected patients are able to opsonize nRBCs, decreasing their deformability and increasing their *in vitro* phagocytic uptake, which may contribute to vivax malaria-associated anemia [5]. However, the relationship between this clearance and ABO blood groups has yet to be investigated.

It has been demonstrated that the ABO blood group system may be associated with severe disease in *P. falciparum* infections, being individuals with A phenotype at a higher risk to develop severe illness while those with type O erythrocytes are more likely protected [6-8]. On the other hand, an association between ABO system and *P. vivax* infection has not been clearly defined and a few studies conducted in this field reveal that hemoglobin levels are lower in blood type O individuals, who are more likely to develop anemia [9, 10]. Taking into account the above-mentioned and our ability to study *in vitro* deformability and clearance of nRBCs [5, 11], we investigated here whether there is an association between ABO blood groups and IgG and IgM autoantibody-driven destruction of nRBCs in *P. vivax* infection.

## 2. Methods

### 2.1. Study population

Blood samples were collected from 139 *P. vivax*-infected individuals attending the Hospital Universitário Júlio Müller, in Cuiabá, Mato Grosso State, Brazil. These subjects were examined by an experienced physician, who applied a questionnaire to collect demographic and epidemiological data. A detailed description of the characteristics of the studied population has been published elsewhere [12]. Blood samples were collected upon admission and analyzed for hemoglobin level. For the current study, anemia was set as hemoglobin levels ≤ 11 g/dl. Parasite species was determined by thick blood smear, as well as by nested PCR amplification of species-specific sequence of the 18S SSU rRNA gene of *Plasmodium* [13]. Based on laboratory results of complete blood count, patients were assigned into two groups: (i) malaria patients with anemia (n = 12) and (ii) malaria patients without anemia (n = 127). A third group included malaria-naïve controls who lived in a non-malariaous area (Belo Horizonte, Minas Gerais state, Brazil) and had never reported exposure to *Plasmodium* parasites (n = 11). Ethical approval was obtained from The Ethics Committee of the National Information System on Research Ethics Involving Human Beings (SISNEP – CAAE: 01496013.8.0000.5149). All individuals included in this study were anonymized, and they provided written informed consent for the collection of samples and subsequent analysis.

### 2.2. ABO typing and erythrocytes isolation

RBC typing was done with commercially available anti-A and anti-B reagents by standard hemagglutination techniques. After typing, nRBCs from healthy volunteers of each blood group, A or O, were isolated from whole blood by Ficoll-Paque Plus density gradient (GE Healthcare, Wauwatosa, WI, USA).

### 2.3. Cell-ELISA

Cell-ELISA was performed according to Mourão et al., 2016. Briefly, 1 × 10^7^ human erythrocytes diluted in PBS containing 1 % (w/v) BSA (PBS/BSA) were added to each well of a 96-well, flat-bottomed, polystyrene microplate (Corning Incorporation, Corning, NY, USA), followed by an overnight incubation at 4°C. Then, erythrocytes were washed five times with PBS and wells were blocked with 5 % BSA for 2 h at 37 °C. Wells were washed again and incubated with sera samples (in duplicate) diluted 1:100 in PBS/BSA for 2 h at 37 °C. Next, wells were incubated with HRP-conjugated anti-human IgG or IgM antibodies (Sigma-Aldrich) diluted 1:500 in PBS/BSA for 90 min at 37 °C. The binding was revealed with 0.5 mg/mL o-phenylenediamine dihydrochloride substrate (OPD) (Sigma-Aldrich, St Louis, MO, USA), and Optical Density (OD) was measured at 492 nm in a Spectra Max 250 microplate reader (Molecular Devices, San Jose, CA).

### 2.4. IgG and IgM purification

For erythrophagocytosis assay and defocusing microscopy we used pooled plasma from each studied group. IgG and IgM antibodies were purified from plasma samples by affinity chromatography using prepacked HiTrap™ Protein G HP columns (GE Healthcare, Wauwatosa, WI, USA) and HiTrap™ IgM Purification HP columns (GE Healthcare, Wauwatosa, WI, USA), respectively, according to the manufacturer’s recommendations.

### 2.5. Defocusing Microscopy

Defocusing Microscopy (DM) is a non-invasive and powerful optical microscopy technique that allows the evaluation of morphological (volume, surface area, sphericity index, total 3-D shape), chemical (hemoglobin content and concentration) and mechanical properties (dynamic membrane fluctuation) of single cells using simple methods and easy setup [11]. DM was used to investigate the effects of different antibodies in the biomechanical properties of erythrocytes through measurement of membrane fluctuations amplitude of individual nRBCs from blood group A and O. DM experiments were performed in a Nikon Eclipse TI inverted microscope as previously described [5, 11]. To determine whether there is a difference between membrane rigidity of O and A erythrocytes after the addition of antibodies purified from different groups of study, we obtained the relative membrane fluctuation, which is calculated as the ratio between membrane fluctuation before and 30 min after antibody addition.

### 2.6. Erythrophagocytosis

Blood samples were collected by venous puncture in heparin tubes from three healthy volunteers from each blood group (A or O) on the day of the erythrophagocytosis assay, which was performed as previously described [5]. Phagocytosed RBCs were counted on 400 macrophages. This count was repeated blinded for three times by each of two microscopists. The erythrophagocytosis rate (ER) was calculated as follows: ER = (phagocytosed RBC/macrophage) x 100.

### 2.7. Statistical analysis

Statistical analysis was performed using GraphPad Prism 5 software. To test if data sets fitted a Gaussian distribution, Kolmogorov-Smirnov test was performed. Data sets that displayed normal distribution were analyzed by Student’s t test (two-tailed) or One-way ANOVA followed by Tukey’s multiple comparison test as appropriate. For nonparametric data, Mann-Whitney rank sum test or Kruskal-Wallis followed by Dunn’s post hoc test was performed as appropriate. p values less than 0.05 were considered significant.

## 3. Results

### 3.1. Anemic patients exhibit higher levels of IgG and IgM antibodies against blood group O erythrocytes than blood group A

We found that anemic patients with acute *P.* vivax malaria have higher IgG levels against erythrocyte molecules from blood group O (median [interquartile range], 1.90 [0.62-3.63]) than to blood group A (0.79 [0.48-2.06]) (p = 0.0161). IgG antibody response among non-anemic patients was also increased to O erythrocytes (1.14 [0.69-1.68]) in comparison to A RBCs (0.69 [0.41-1.24]) (p < 0.0001) (Fig. 1A).

**Fig. 1.**
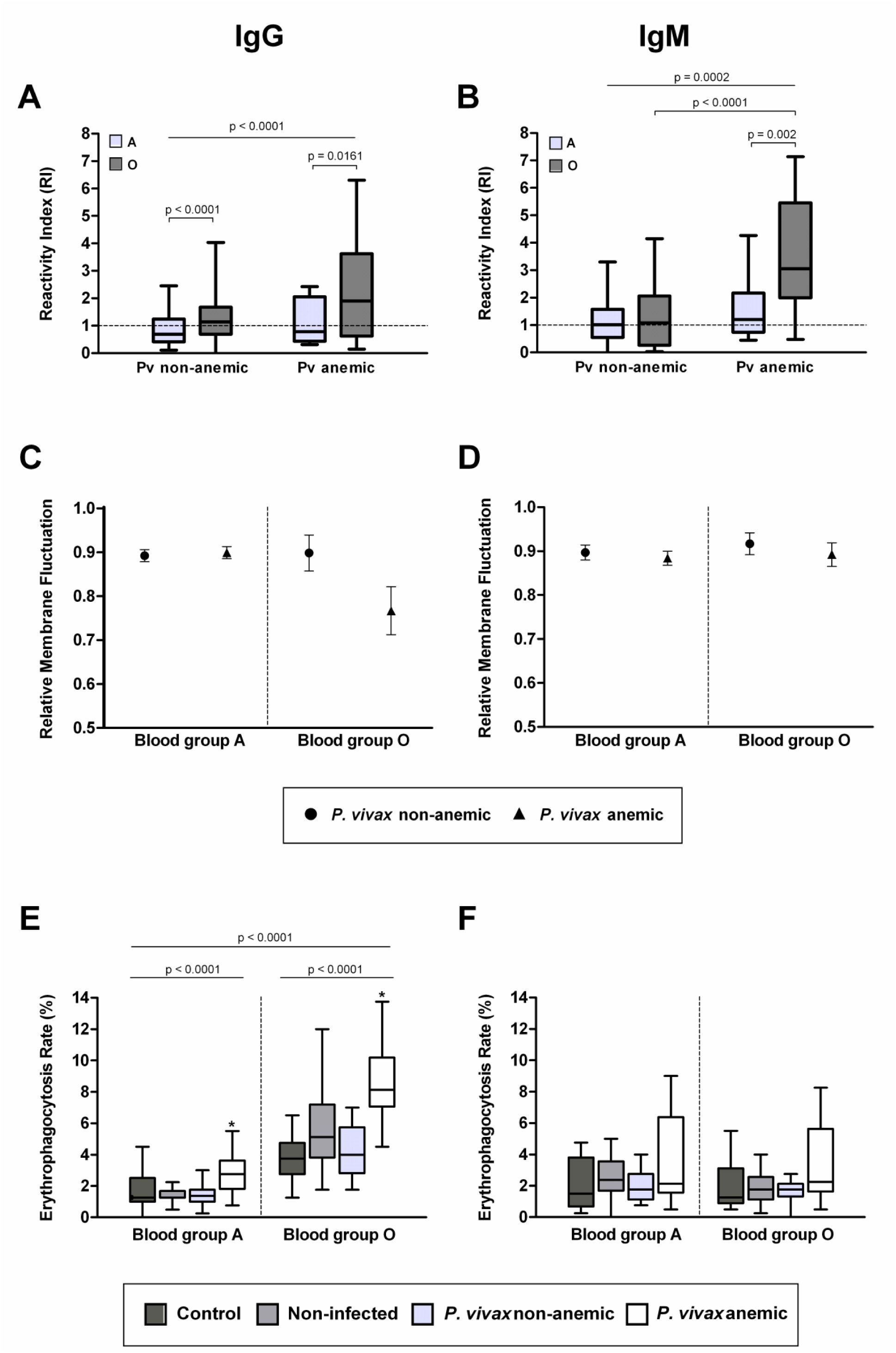
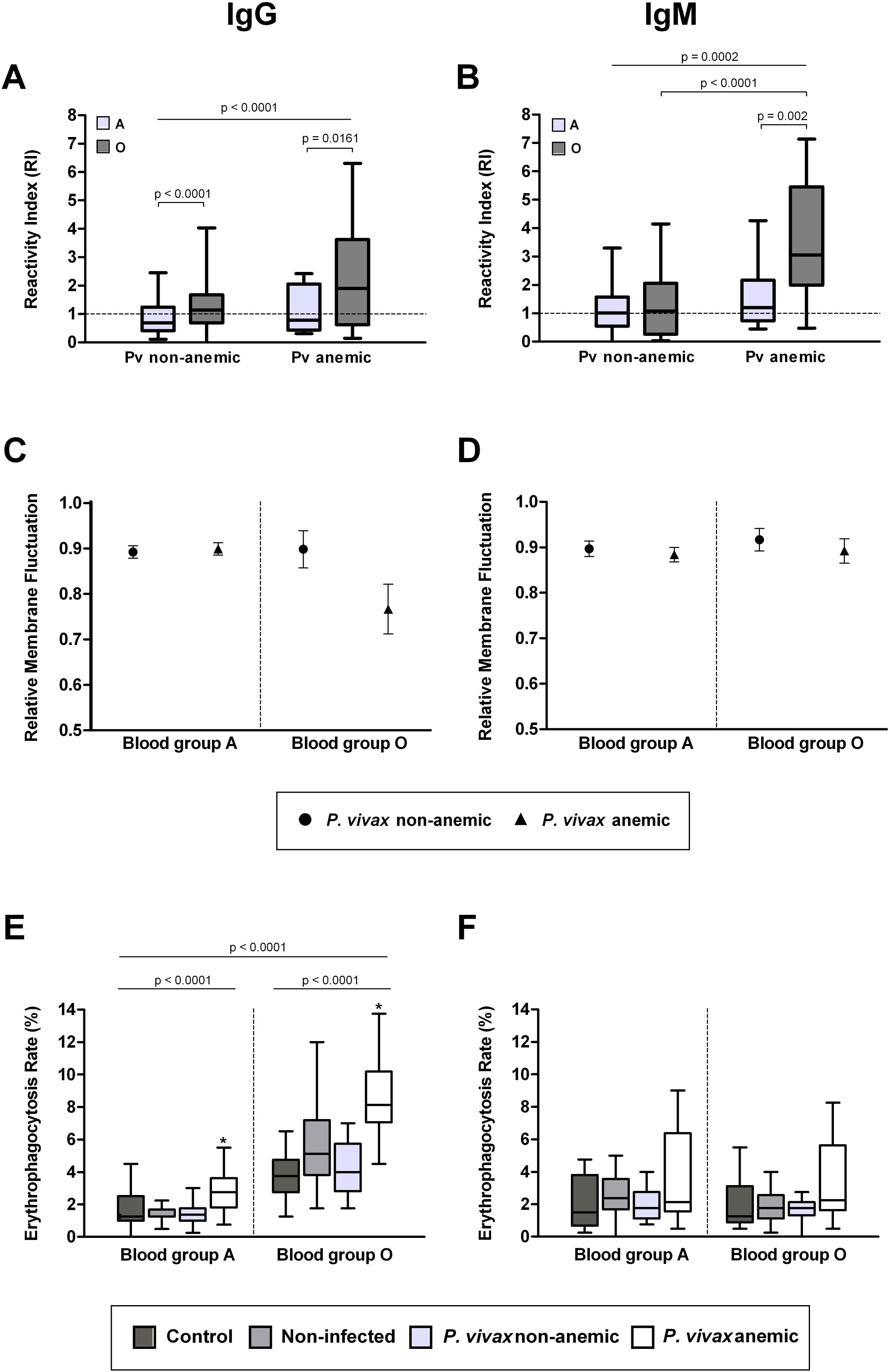
Effects of anti-RBC IgG and IgM from patients with acute vivax malaria on erythrocytes from blood group A and O. Levels of IgG **(A)** and IgM **(B)** against nRBCs were detected in plasma from non-anemic (n = 127) or anemic *P. vivax*-infected patients (n = 12) by cell-ELISA. Antibody responses were expressed as reactivity index (RI) which was calculated as the ratio between the mean OD from each sample duplicate and the mean OD plus two standard deviations of samples from 11 malaria-naïve blood donors never exposed to malaria. RIs equal or greater than 1.0 were scored as positive and smaller as negative. *P* values < 0.05 were considered significant. Data are presented as box-and-whiskers plots with interquartile ranges. Horizontal lines indicate median. Modifications in the amplitude of membrane fluctuations of single human RBCs from blood group A and O obtained from healthy donors were examined by defocusing microscopy before and 30 min after the addition of IgG **(C)** and IgM **(D)** purified from anemic (n = 7) and non-anemic (n = 7) *P. vivax* patients. RBC membrane rigidity was expressed as relative membrane fluctuation that is represented by the ratio between membrane fluctuation values before and after antibody addition. The symbols indicate the mean values over 10 nRBCs and error bars represent the standard deviation. To assess the ability of IgG **(E)** and IgM **(F)** to induce erythrophagocytosis, erythrocytes from blood type A (n = 7) and O (n = 7) were collected from healthy donors and opsonized with mentioned purified pooled antibodies from non-infected individuals, non-anemic and anemic *P. vivax* patients. As control, PBS was used instead of immunoglobulins. The erythrophagocytosis rate was expressed as ER and calculated as follows: ER = (phagocytosed RBC/macrophage) x 100. The results of three independent experiments for each blood type and antibody are presented as box-and-whiskers plots with interquartile ranges. Medians are indicated by horizontal lines. *P* values < 0.05 were considered significant.

In addition, anemic patients with patent *P. vivax* infection exhibited higher levels of IgM against erythrocytes from blood group O (3.06 [2.00-5.46]) than to blood group A (1.20 [0.74-2.17]) (p = 0.002) (Fig. 1B).

### 3.2. O erythrocytes coated with IgG from anemic-*P. vivax*-infected patients exhibit a higher increase in membrane rigidity in comparison to A erythrocytes

Since changes in erythrocyte membrane fluctuations indicate alterations in the biomechanical properties of RBCs, we investigated by defocusing microscopy the effects of IgG and IgM antibodies purified from anemic and non-anemic *P. vivax*-infected patients on O and A erythrocytes. Our results showed that membrane fluctuations of blood group A erythrocytes exhibited a reduction of 10% (before: 17.15 ± 3.48 nm, after: 15.46 ± 3.40 nm; p = 0.0001) after the addition of IgG from anemic patients and 11% for non-anemic (before: 15.06 ± 1.84 nm, after: 13.40 ± 1.43 nm; p = 0.0001). For the group O, we observed a decline of 21% for anemic (before: 24.02 ± 5.15 nm, after: 18.95 ± 7.34 nm; p = 0.0007), *versus* 11% for non-anemic ones (before: 25.50 ± 3.35 nm, after: 22.66 ± 2.80 nm; p = 0.0331) (Fig. 1C). For type A and O nRBCs, a decrease on membrane fluctuation was also observed when these cells were coated with IgM from non-anemic and anemic *P. vivax*-infected patients (Fig. 1D). The mean value of membrane fluctuation for group A decreased 12% after the addition of IgM from non-anemic patients (before: 17.34 ± 2.13 nm, after: 15.61 ± 2.42 nm; p = 0.0001), while a decrease of 10% was found for anemic ones (before: 20.34 ± 4.00 nm, after: 17.87 ± 3.00 nm; p = 0.0001). In the case of group O erythrocytes, the mean values of membrane fluctuation decreased 10% after addition of IgM from both anemic (before: 25.38 ± 4.30 nm, after: 22.69 ± 4.68 nm; p = 0.0029) and non-anemic patients (before: 24.98 ± 3.54 nm, after: 22.95 ± 4.16 nm; p = 0.0003). In summary, these results indicate that erythrocytes from blood group O coated with IgG from anemic *P. vivax*-infected patients exhibit a higher decrease in membrane fluctuation. Despite a decrease on membrane fluctuation has been observed after addition of IgM, no difference on the degrees of reduction between A and O erythrocytes were found, as opposed to IgG. Finally, no changes in RBC shape were observed after addition of different antibodies.

### *3.3. In vitro* phagocytosis is higher in blood group O than in A erythrocytes coated with IgG from anemic patients with acute *P. vivax* malaria

Because IgG and IgM of *P. vivax*-infected patients were able to decrease nRBC membrane deformability in both A and O erythrocytes, even in different degrees, we used an *in vitro* system to evaluate their aftermath in the phagocytic activity of a human monocytic cell line. Fig. 1E and Fig. 1F show median rates of erythrophagocytosis observed for A and O erythrocytes opsonized with purified immunoglobulins from healthy donors, anemic or non-anemic *P. vivax*-infected patients. Our results indicate that IgG purified from anemic *P. vivax*-infected patients increases *in vitro* phagocytosis of both A (median [interquartile range], 2.75 [1.81-3.63]) and O erythrocytes (8.125 [7.06-10.19]) in comparison to control (A: 1.23 [1.00-2.50]; O: 3.75 [2.75-4.75]), non-infected (A: 1.25 [1.25-1.69]; O: 5.13 [3.81-7.19]) and non-anemic patients (A: 1.38 [1.00-1.75]; O: 4.00 [2.81-5.75]). These data highlight an increased susceptibility of O nRBCs to *in vitro* phagocytosis (Fig. 1E).

In relation to IgM (Fig. 1F), no significant differences were detected among nRBCs from different blood groups and either among antibodies from the different studied groups. These results indicate that although IgM deposited on nRBCs decreases cell membrane fluctuation, it plays no role on clearance by phagocytosis.

## 4. Discussion

This is one of the first studies that report a possible association between ABO blood groups and the potential removal of nRBCs mediated by autoantibodies in *P. vivax* malaria. Results reported here suggest that: (1) *P. vivax*-infected patients with anemia have increased levels of IgG and IgM against nRBCs, mainly of those from O blood type in comparison to non-anemic ones; (2) IgG and IgM purified from anemic patients are able to increase nRBCs membrane rigidity, an effect that is enhanced in O erythrocytes coated with IgG; (3) IgG but not IgM promote *in vitro* phagocytosis of nRBCs and the phagocytic uptake is increased in erythrocytes from blood group O.

Our results are in accordance with a few previous studies [9, 10]. Resende et al. genotyped five single nucleotide polymorphisms at the ABO gene of Brazilian Amazon patients and associated this information to hematological data, showing a relationship between blood group O and increased susceptibility to anemia in *P. vivax* infection [9]. A similar result was also found by Kuesap & Na-Bangchang that associated parasitemia and blood parameters indicative of anemia to ABO blood group in patients along Thailand and Myanmar [10]. Besides finding similar results, here we provide additional information highlighting *in vitro* mechanistic evidence in favor of a higher susceptibility to anemia in *P. vivax*-infected patients from blood group O.

On the other hand, a systematic review and meta-analysis using data from *P. falciparum* infections revealed that individuals with O blood group are less susceptible to severe malaria [8]. Indeed, it has been postulated that the resistance conferred by blood group O during falciparum malaria may be due to the lower percentage of infected RBCs forming rosettes, which are smaller and more fragile in comparison to non-O blood groups [6, 14]. In addition, it has also been demonstrated that infected O erythrocytes are more phagocytosed by macrophages than non-O RBCs. This enhanced clearance could be due to an increased hemichrome deposition and band 3 aggregation in infected O erythrocytes, suggesting that it may confer protection against severe *P. falciparum* malaria [15]. However, this protective effect of blood group O observed in falciparum malaria has been recently countered by Theron et al. who demonstrated that *P. falciparum* preferentially invades O erythrocytes *in vitro* [16]. It is worth mentioning that there are differences between *in vivo* and *in vitro* assays, and both are important to define the mechanisms of protection.

It has long been known that immune response, life cycle and disease pathogenesis are different for both *P. vivax* and *P. falciparum*, parasites which have their own peculiarities. Thus, it is not surprising that ABO blood groups influence the outcome of *Plasmodium* infection in different manners.

Another point that should also be taking into account is the fact that O erythrocytes express only the H antigen on their surface, making it unlikely to anti-A and anti-B antibodies to bind to this antigen. Therefore, the higher antibody response observed against O erythrocytes in the current study may not be due to naturally acquired anti-ABO antibodies, but to a parasite-driven immune response.

Our defocusing microscopy and erythrophagocytosis results indicate that IgG antibodies purified from anemic patients with vivax malaria can decrease nRBCs membrane dynamics fluctuations and increase their *in vitro* phagocytic uptake, mainly of those from O phenotype. This clearance of non-infected erythrocytes is an important component of malaria-associated anemia as previously demonstrated [5, 17] and suggests that O individuals may be more prone to develop anemia during *P. vivax* infections. These data add new insights about the possible association between ABO blood groups and *P. vivax*-associated anemia. Further investigation should be carried out to assess why O erythrocytes are more susceptible to clearance.

Another result that is worth mentioning is the detection of high levels of IgM antibodies against nRBCs in *P. vivax*-infected patients. It is possible that those antibodies may recognize phosphatidylserine, which is exposed in the surface of both infected and non-infected RBCs during malaria [17], as it has been demonstrated by Barber et al. [18]. Further studies are necessary to elucidate this hypothesis.

The increased IgM response, in addition to its low ability to stimulate *in vitro* erythrophagocytosis, suggest that those antibodies may play an alternative role in nRBC destruction, especially of O erythrocytes. We believe that IgM may exert a pathogenic role in *P. vivax*-associated anemia through multivalency-dependent hemagglutination as it has already been demonstrated in a mouse model of autoimmune hemolytic anemia [19], an issue that also warrants further investigation.

In conclusion, the findings reported here give new information about the immune response associated with clearance of nRBCs as well as its relationship with ABO blood groups in *P. vivax* infection. In order to fully understand what makes an anti-RBC autoantibody pathogenic, further studies are essential.

## Acknowledgements

The authors would like to thank all patients and their families who contributed to the current study. We are also grateful to health professionals and students from UFMT.

## Financial support

This work was supported by Conselho Nacional de Desenvolvimento Cientifico e Tecnológico [grant numbers: 309202/2013-2; 158045/2015-7; 404365/2016-7; 154378/2018-6]; Fundação de Desenvolvimento da Pesquisa do Estado de Minas Gerais [grant number: APQ000361-16]. The funders had no role in study design, data collection and analysis, decision to publish, or preparation of the manuscript.

## Conflict of interest

The authors declare no conflict of interests, and this manuscript has not been accepted or published elsewhere.

